# Sequence models conditioned on splicing factor expression predict splicing in unseen tissues

**DOI:** 10.64898/2026.01.20.700496

**Authors:** Aniketh Janardhan Reddy, Peter H. Sudmant, Nilah M. Ioannidis

**Author notes:** These authors jointly supervised this work (Peter H. Sudmant, Nilah M. Ioannidis).

## Abstract

Predicting how RNA splicing varies across tissues is important for understanding the impact of genetic variation and identifying splicing-based disease mechanisms. Although many sequence-based deep learning models have been developed to predict splicing, most predict splice sites rather than full splicing events, are restricted to tissues seen during training, or do not account for trans-regulatory variation such as differences in splicing factor expression. Here, we present Splice Ninja, a sequence-based deep learning model that predicts percent spliced-in (PSI) values for individual splicing events across tissues by conditioning on the expression levels of 301 splicing factors. Trained on PSI measurements from many different human tissues and cell types, Splice Ninja is evaluated on three entirely held-out tissues. Despite not seeing these tissues during training, it makes accurate PSI predictions and can identify a substantial fraction of splicing events with high tissue-specificity. Its performance is comparable to Pangolin [1], which is trained directly on the test tissues, but falls short of TrASPr [2], a substantially larger model also trained on the test tissues. Splice Ninja demonstrates that integrating trans-regulatory context into sequence-based splicing models enables generalization to new cellular environments. This framework offers a promising direction for building robust, context-aware predictors of alternative splicing. Our code is available at https://github.com/anikethjr/splice_ninja.

## 1 Introduction

Alternative splicing (AS) enables a single gene to produce multiple mRNA and protein isoforms, expanding the functional diversity of the human transcriptome. More than 95% of multi-exon human genes undergo AS [3], and precise control of isoform expression is essential for normal cellular function. Disruptions in splicing regulation contribute to a wide range of diseases, including cancer and neurodevelopmental disorders [4]. Mutations can alter splicing either through cis effects, by disrupting binding sites for RNA-binding proteins (RBPs) that regulate splicing, or through trans-effects, by perturbing the expression levels of either RBPs or other splicing-associated proteins. More generally, proteins that are involved in the core splicing machinery or in regulating splicing are called splicing factors (SFs). Although several sequence-based models have been developed to predict splicing outcomes, major challenges remain. Most models either do not make tissue-specific predictions at all [5, 6] or can only make predictions for tissues seen during training [1, 7, 8]. They typically predict individual splice sites and their usage levels rather than full splicing events [1, 5], and they fail to account for variation in SF expression, limiting their ability to model trans-effects [1, 5–8].

To overcome these limitations, an ideal splicing predictor should satisfy three key desiderata: (1) it should make predictions in unseen tissues or conditions, enabling generalization beyond the tissues seen during training; (2) it should model full splicing events—not just individual splice sites—to provide more interpretable predictions; and (3) it should incorporate SF expression to account for trans-regulatory effects. A model that satisfies these criteria would have broad utility across genomics and disease biology. For instance, it could predict the effects of an individual’s splicing-disruptive mutations in tissues that are difficult to sample, such as the brain, by leveraging population-level SF expression data from these tissues. In single-cell RNA-seq datasets, where splicing quantification is limited due to low read depth or the use of techniques such as 5’-cap capturing, such a model could estimate splicing patterns using more readily available SF expression profiles. Moreover, by conditioning on SF expression, the model can capture the effects of trans-acting variants that alter the splicing machinery itself—an important class of regulatory variants that are otherwise difficult to model.

In this work, we introduce Splice Ninja, a deep learning model that fulfills all three desiderata outlined above. Splice Ninja makes tissue-specific predictions of splicing events by combining sequence information with the expression levels of splicing factors in the target tissue. It predicts percent spliced-in (PSI) values, a widely used metric that quantifies the proportion of transcripts incorporating a given alternate exon segment. The model is trained on a large dataset of splicing events measured across ~150 human tissues and cell types from VastDB [9], and evaluated on three held-out tissues that are entirely excluded during training. Our results show that Splice Ninja makes accurate PSI predictions for most splicing events in these unseen tissues and successfully recovers many tissue-specific splicing changes—despite the inherent difficulty of this task. When compared to existing methods trained directly on the test tissues, Splice Ninja performs comparably to Pangolin [1], a model with similar architecture and scale, though it lags behind TrASPr [2], a substantially larger model. These findings highlight the potential of models that integrate trans-regulatory context via SF expression: even without direct exposure to a target tissue during training, Splice Ninja approaches the performance of models that were directly trained on the target tissue.

## 2 Related work

Understanding the regulation of splicing and building an in-silico “splicing code” that can take in a genomic sequence, annotate introns and exons, and/or predict the levels of different isoforms while considering cellular conditions, has long been a central goal in computational biology. Such models are essential for interpreting the functional effects of variants and developing splicing-targeted therapies. In this section, we review both classical models that rely on hand-engineered features and modern deep learning approaches that directly model sequence. We highlight how Splice Ninja builds on these foundations by integrating both sequence and SF expression levels that provide trans-regulatory context.

### 2.1 Feature-based models

Early work focused on recognizing 5’ and 3’ splice sites, and regulatory elements that control splicing using relatively simple machine learning or statistical models. For example, the popular MaxEntScan method [10] used maximum entropy modelling to detect 5’ and 3’ splice sites. Then, to study AS, Barash et al. [11] presented a model that can predict cassette-exon usage in different mouse tissues using various motif and transcript structure-based features. Zhang et al. [12] extended this framework by incorporating thousands of exon sequence features and RBP expression levels into a neural network that predicted if there would be differential splicing between two biological conditions (e.g. cell types). While modern deep learning models have largely moved beyond hand-crafted features, most do not model trans-effects mediated by SF expression. Splice Ninja addresses this gap by combining sequence-based deep learning with explicit modelling of SF expression. TrASPr [2], a contemporary sequence-based deep learning model, also utilizes RBP expression levels to capture trans-effects, but their RBP-conditioned models are not publicly available.

### 2.2 Sequence-based deep learning models

Most modern splicing predictors fall into two broad categories: (1) models that predict individual splice sites, and (2) models that predict full-event inclusion levels, typically quantified as PSI.

SpliceAI [5] was one of the first deep learning models to predict splicing directly from sequence, using a deep residual network (ResNet) [13] to identify annotated splice sites in a context-agnostic manner. SpliceAI is more accurate than traditional approaches like MaxEntScan for recognizing splice sites, and it can help identify splicing-altering pathogenic variants. Pangolin [1] extended SpliceAI to predict tissue-specific splicing by jointly training on data from multiple tissues and species using multi-task learning (MTL). In addition to predicting the occurrence of splice sites in each tissue seen during training, Pangolin also predicts tissue-specific splice site usage levels. More recent work by You et al. [8] introduced a hybrid convolutional neural network (CNN) and transformer-based architecture for tissue-specific splice site prediction, while Chen et al. [14] and de Almeida et al. [15] fine-tuned large-scale genomic language models for similar tasks.

The second class of models directly predict PSI values. MMSplice [6] predicts the impact of sequence variants on splicing using modular neural networks trained on GENCODE annotations and PSI values from MPRA data. MTSplice [7] builds on this to enable tissue-specific PSI prediction in seen tissues using MTL. TrASPr [2] is a more recent BERT-based model [16], pretrained to predict splice site annotations and fine-tuned to predict tissue-specific PSI for exon skipping events. TrASPr is trained on six GTEx tissues [17] and takes in sequences around splice junctions along with exon/intron lengths, conservation scores, and a tissue pair. It is trained to predict the PSI value in the first tissue and the PSI difference between the two tissues (dPSI). TrASPr outperforms methods like SpliceAI and Pangolin (after converting their outputs to PSI estimates) for exon skipping PSI prediction, including for tissue-specific events. Like Splice Ninja, TrASPr can also make predictions for unseen tissues when trained using principal components of RBP expression levels as additional inputs.

A third class of models that directly predict RNA-Seq coverage is emerging. Borzoi [18] is an example of such a model. It uses a UNet-style architecture [19] in conjunction with transformers, and is trained using data from many RNA-Seq experiments and other datasets that measure transcription factor-binding, accessibility, expression, and histone modifications along the genome. However, Borzoi also cannot make predictions for unseen tissues, does not incorporate trans-effects, and computing splicing event-level metrics, such as PSI, from RNA-Seq coverage predictions is difficult. AlphaGenome [20] builds on Borzoi to jointly predict RNA-seq coverage, splice site usage, and junction usage, enabling event-level splicing metric computation; however, like Borzoi, it cannot generalize to unseen tissues.

In this work, we adopt the tissue-specific PSI prediction paradigm, which allows for modelling full splicing events. However, unlike many existing models, Splice Ninja is designed to generalize to tissues unseen during training by conditioning predictions on measured SF expression levels. We compare Splice Ninja’s performance on unseen tissues to the performance of SpliceAI, Pangolin, and TrASPr using some of the benchmarks presented in the TrASPr manuscript. Crucially, data from the tissues we held out for testing were used to train both Pangolin and TrASPr. Our results show that Splice Ninja achieves comparable performance to Pangolin without requiring training data from the target tissues. Although the TrASPr manuscript has some results illustrating TrASPr’s performance on unseen GTEx tissues, at the time of writing, we could not find the pretrained model or data used for their unseen tissue analyses. Therefore, we do not directly compare the performance of Splice Ninja and TrASPr on tissues unseen by both models at this time.

## 3 Problem statement

In this section, we formally define the splicing prediction task addressed in this work. We focus on four common types of alternative splicing events: exon skipping (ES), intron retention (IR), alternative acceptor (ALTA), and alternative donor (ALTD), as illustrated in Figure 1. In any given RNA-Seq experiment, and for a given splicing event, percent spliced in or PSI values quantify the proportion of total transcripts that contain the alternate exon segment^1^. We use PSI values computed using the vast-tools package [9].

**Fig. 1:**
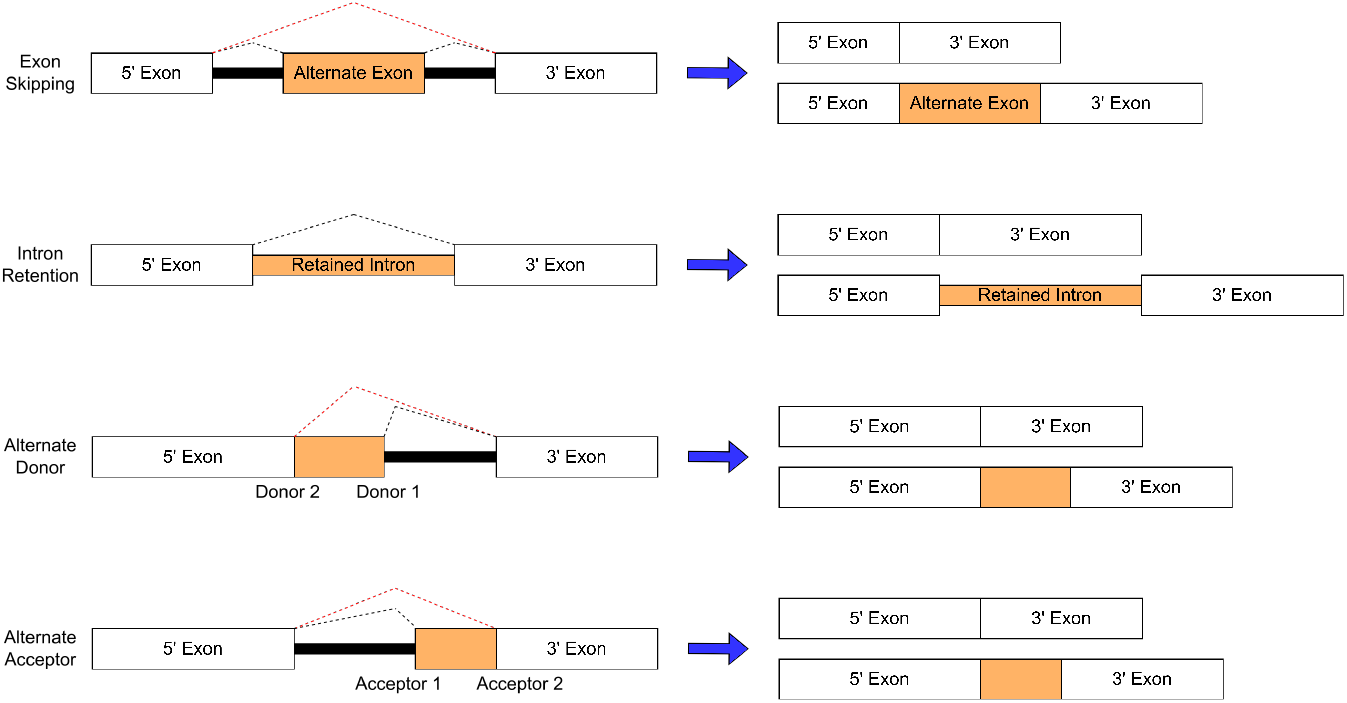
The four main types of splicing events that we focus on in our work. Here, each block represents an exon and the solid lines between them represent introns. The black dashed lines indicate the exon-exon junctions used in the first transcript and the red dashed lines indicate the junctions used in the second one. The alternate exon segments are shown in orange and PSI measures the ratio of the number of transcripts that contain this segment to the total number of transcripts.

Let 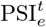 denote the observed PSI value for a splicing event *e* in tissue *t*. We are given a training set consisting of PSI measurements for a set of *m* splicing events *e*_1_, …, *e*_*m*_ across *n* tissues *t*_1_, …, *t*_*n*_. Importantly, these events are drawn from a set of training chromosomes that are disjoint from those used in the validation and test sets. For each tissue *t*, we are also provided with the expression levels of 301 SFs, denoted by the vector *sf*^*t*^ ∈ ℝ^301^.

The goal is to predict the PSI value for an event *e*′ in a tissue *t*′ that was not observed during training. Specifically, we aim to learn a model:

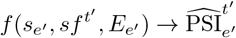

that takes as input:

1. the one-hot encoded 10,000 bp genomic sequence *s*_*e*_′ containing the splicing event and surrounding context,
2. the SF expression vector 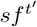 for the target tissue,
3. and the one-hot encoded event type *E*_*e*_′ ∈ {ES, IR, ALTA, ALTD}, and outputs the predicted PSI value 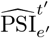.

## 4 Predicting tissue-specific splicing with Splice Ninja

We introduce Splice Ninja, a sequence-based deep learning model designed to predict tissue-specific splicing patterns, including for tissues unseen during training. We first describe its architecture before detailing how it is trained using data from VastDB [9]. Finally, we describe how a trained model can be used for predicting splicing in unseen tissues.

### 4.1 Model architecture

Figure 2 illustrates Splice Ninja, with Figure S1 providing more details about the various components. Its architecture is based on the largest SpliceAI model that takes in 10,000 bp sequences as input [5], with modifications to facilitate tissue-specific PSI predictions.

**Fig. 2:**
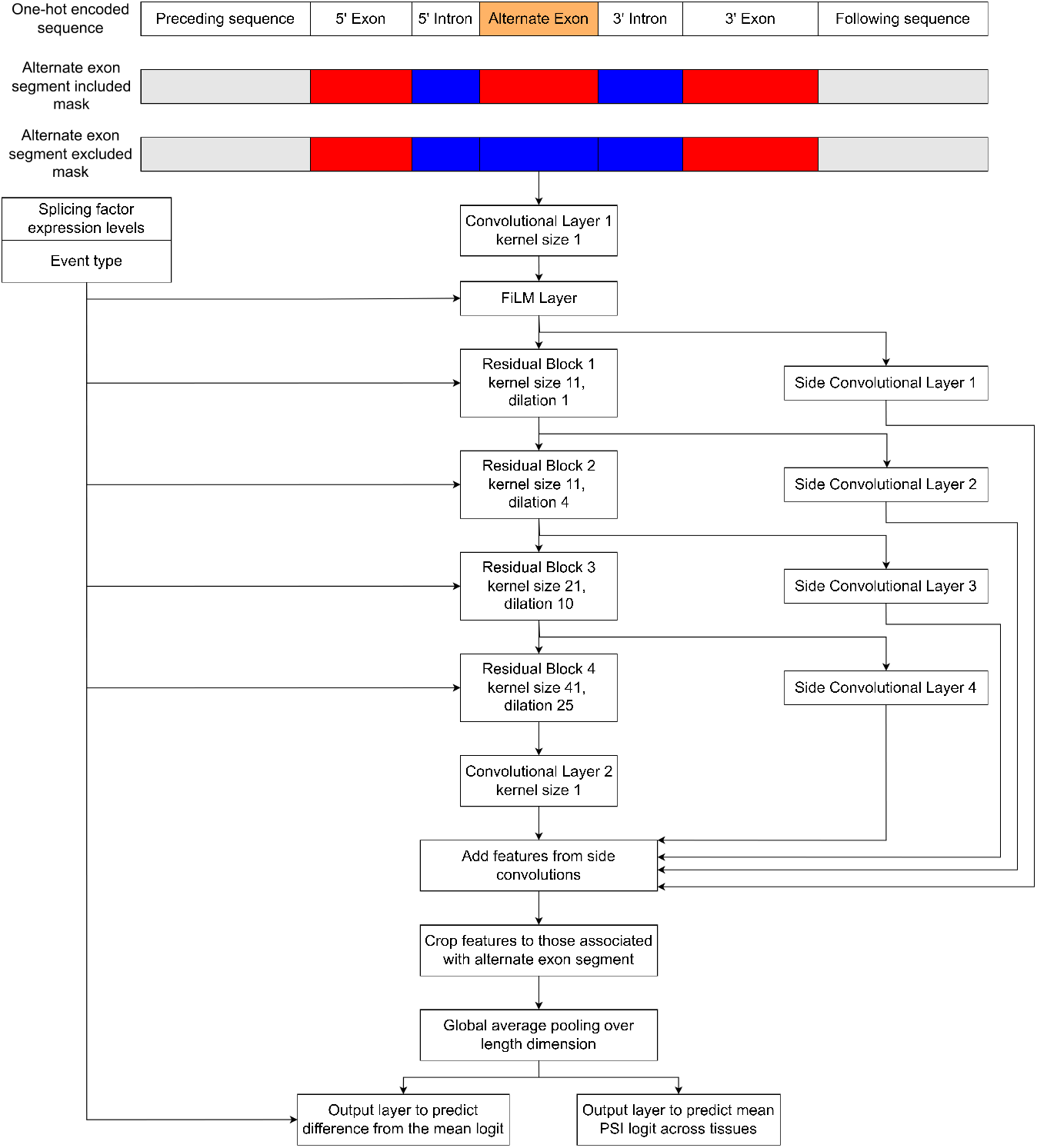
Splice Ninja model to predict tissue-specific PSI values using sequence and SF expression levels. In the masks, the red blocks indicate exons (assigned a value of 1), the blue blocks indicate introns (value −1), and the grey blocks indicate surrounding regions (value 0). An exon skipping event is shown here but the same modelling scheme can be used for any splicing event type.

Splice Ninja takes in a 10,000 bp genomic sequence encompassing the splicing event and its surrounding context^2^. It also takes in two masks that define the splicing event for which PSI predictions are to be made. These masks indicate exon and intron structures when the alternate exon is either included or excluded (Figure 2). The one-hot encoded genomic sequence and masks are concatenated and processed by a ResNet backbone [13].

In order to make tissue-specific predictions, Splice Ninja also takes in SF expression levels in a given tissue, allowing it to infer the regulatory environment of the tissue and incorporate this information while making predictions. Rogalska et al. [21] assembled a comprehensive set of 305 SFs and performed knockdown experiments to assess their roles. We choose a subset of 301 SFs for which expression information is available in VastDB [9], the dataset we use in our experiments. After computing the logarithm of the transcripts per million (log_2_ TPM) for each SF, we normalize them across the 301 SFs so that they sum to 1. This 301-dimensional vector captures the relative levels of different SFs in different tissues. Since we jointly train on all splicing event types, we also supply a one-hot encoded vector that specifies the event type (one of ES, IR, ALTA, or ALTD). Together, the SF expression and event type vectors represent the condition information used to make PSI predictions. This condition information is used throughout the model in Feature-wise Linear Modulation (FiLM) layers [22] that compute shift and scale vectors. Given a convolutional kernel *k*’s activation 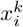 at position *i* of the input, the FiLM layer computes a learned affine transformation 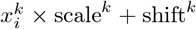 that adjusts the activation based on the condition information. Figure S1 provides more details on the FiLM layer.

Rather than directly predicting PSI values, Splice Ninja separately predicts the mean PSI logit across tissues and the tissue-specific deviation from this mean. Adding these two logits and applying the sigmoid function gives us tissue-specific PSI predictions. We found this prediction scheme to be better than directly predicting PSI values in our experiments. We operate in the logit-space since PSI values across samples are often modelled using the logit normal distribution [23], and using logits leads to improved training stability and gradient behavior.

### 4.2 Model training and data splits

We train Splice Ninja using human data from VastDB [9], which provides uniformly processed PSI and expression measurements across multiple tissues and cell types. We partition the 145 samples into training, validation, and test sets, explicitly reserving cerebellum, liver, and heart samples for the test set to facilitate comparisons with existing methods (sample names listed in Appendix A). Simultaneously, we also divide the chromosomes among the sets. Like SpliceAI [5], Pangolin [1], and TrASPr [2], we use data from chromosomes 1, 3, 5, 7, and 9 in the test set; chromosomes 8, 10, 17, and 22 are used for validation; and all other chromosomes are used in the train set. Additionally, we filter out low quality PSI measurements and any events that are not observed in at least 10 samples to ensure that we train on high quality data. To get SF expression levels, we estimate TPM from the read count data available on VastDB^3^.

In every training batch, we sample from 32 different splicing events and for each event, we sample 12 tissue-specific PSI values (total batch size of 32 × 12 = 384). For most splicing events, we observed that PSI values rarely deviated significantly from the mean PSI value across samples. Here, significant deviation is defined as |dPSI| ≥ 0.15. Therefore, we upsample these significant deviations to allow the model to learn tissue-specific splicing more effectively, making up two-thirds of the PSI values in a batch, with the other one-third being sampled from the remaining values. Then, we use the sum of three losses to train Splice Ninja:

1. Binary cross-entropy (BCE) between the actual and predicted mean PSI
2. Binary cross-entropy (BCE) between the actual and predicted tissue-specific PSI
3. A ranking loss (RL) that forces the model to correctly order samples by PSI for a given event. To compute this loss, we consider every pair *a, b* of PSI values from a given event that are significantly different (|*a* − *b*| ≥ 0.15). If *a* > *b*, this pair is assigned a label *y* = 1; and if *a* < *b, y* = −1. Then, if the predictions are 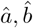, the 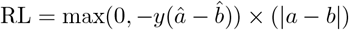. The RL is zero if the pair is ordered correctly and is greater than zero if the ordering is incorrect. We also have a weight term | *a* − *b* | that prioritizes pairs with larger dPSI. The RL losses between all valid pairs are averaged to get the final loss.

We train using the AdamW optimizer [24] with a learning rate of 1e-4 and a weight decay of 1e-4. The model is trained for 25 epochs^4^ using 4 Nvidia A40 GPUs and the checkpoint with the best Spearman correlation on the validation set comprising unseen tissues and chromosomes (epoch 21) is used in all analyses.

### 4.3 Predictions for unseen tissues or conditions

A trained Splice Ninja model predicts PSI values in novel tissues or conditions using their SF expression profiles. As mentioned previously, these expression levels need to be log_2_ TPM values that are normalized to sum to 1. This modelling paradigm allows us to capture trans-effects and is especially beneficial for:

1. Predicting splicing in tissues that are hard to access for RNA-Seq sample collection (e.g. brain). Here, to probe the effects of an individual’s mutations, we could use mean SF expression levels from a population as an input along with the individual’s genome sequence.
2. Predicting splicing at single-cell-resolution. Due to limited read-depth in many single-cell experiments, PSI measurements may be inaccurate due to a limited number of reads that span alternate junctions - these reads are crucial for accurate PSI estimation. However, expression measurements are likely more accurate, enabling us to measure SF expression and thus predict splicing at single-cell-resolution. Moreover, the usage of 5’-cap capturing, a popular single-cell sequencing method, does not allow us to estimate PSI values for events that are located in the middle of the transcript. Using Splice Ninja in such settings could again enable single-cell-resolution splicing predictions.
3. Assessing splicing impacts of trans-acting mutations affecting SF expression, enabling precise identification of mutation-driven splicing changes.

In our analyses, we predict splicing in cerebellum, liver, and heart samples in this manner and show that we are able to make accurate predictions while capturing some tissue-specific variations.

## 5 Results

We demonstrate the utility of Splice Ninja by showing that it can accurately predict splicing events in unseen tissues. Additionally, we compare its performance on these unseen tissues with existing methods that are trained on these tissues and show that its performance is comparable to Pangolin [1] while still trailing TrASPr [2]. This is a notable result since Splice Ninja never observes splicing data from these tissues while Pangolin and TrASPr are directly trained on this data.

### 5.1 Predicting splicing events in unseen tissues

We evaluated Splice Ninja on the test set described in Section 4.2 that comprises data from cerebellum, liver, and heart samples - tissues for which splicing data is never observed during training. Figure 3 illustrates Splice Ninja’s overall prediction performance. For ES, ALTA, and ALTD events, we observe a very high correlation between predicted and measured PSI values, and more than 92% of PSI predictions are within 0.2 of the measurements. However, performance on IR events is much lower, likely due to the narrower distribution of PSI values that might lead to IR events contributing lesser to the overall loss that emphasizes events with large variance in PSI values across samples. In these unseen tissues, we also see that performance on events from train chromosomes is substantially higher than events from test chromosomes. This tells us that observing tissue-specific PSI variations in other tissues allows us to make more accurate predictions in unseen tissues. Thus, a future model trained on events from all chromosomes could enable even better performance for unseen tissues.

**Fig. 3:**
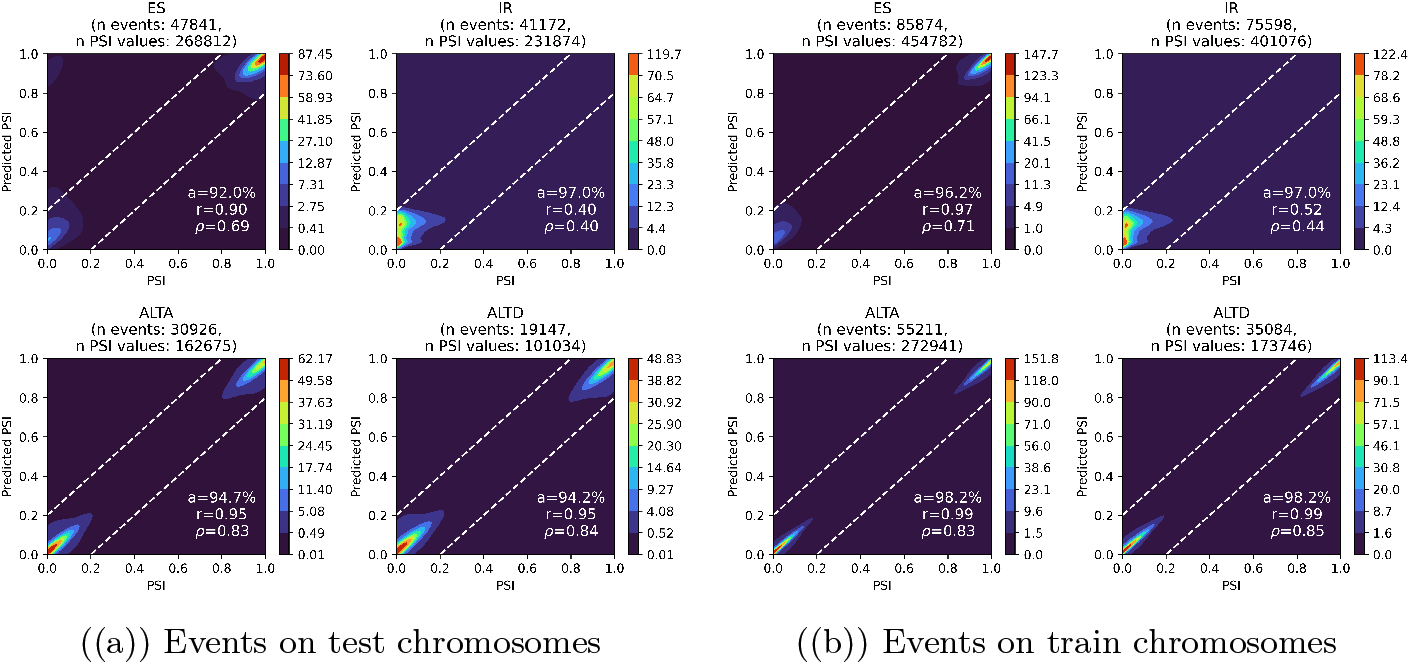
Distributions of measured and predicted PSI values for each splicing event type in unseen tissues. In (a) we evaluate on held-out test chromosomes, and in (b) on unseen tissues with training chromosomes. For each panel, we report the percentage of predictions within 0.2 of the measured PSI (“a”), the Pearson correlation (*r*), and the Spearman correlation (*ρ*) between predicted and measured values. Dashed lines show the ±0.2 deviation boundaries from the identity line.

To better understand Splice Ninja’s ability to make accurate tissue-level predictions, we evaluated prediction performance for each tissue individually. Table 1 presents these results. We observe the same the trends as in Figure 3, with more accurate predictions for ES, ALTA, and ALTD events, and for events from train chromosomes. In addition, performance on all three tissues is comparable across event types.

**Table 1:**
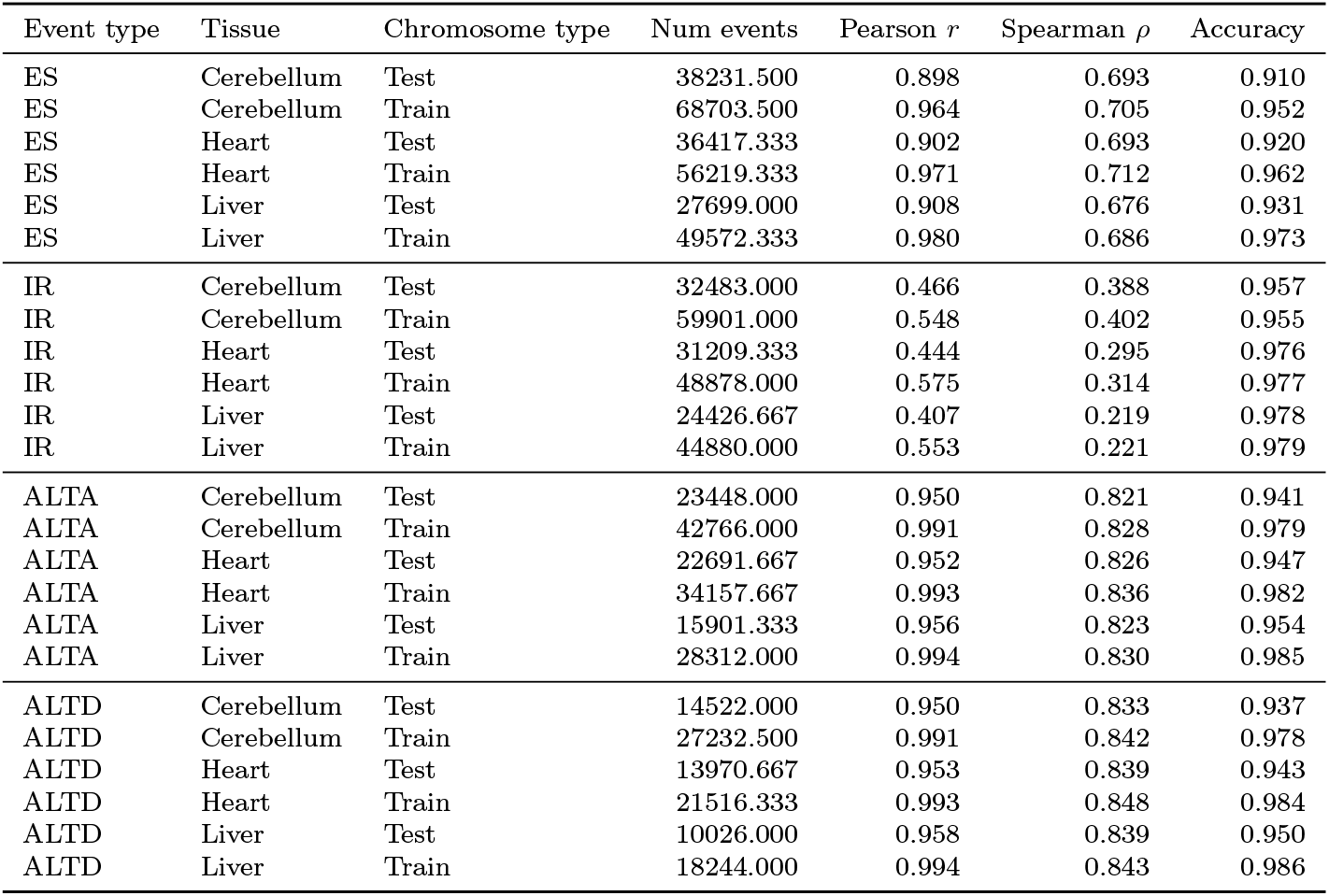
Tissue-level PSI prediction performance for unseen tissues. Metrics are averaged over samples collected from a given tissue (cerebellum has 2 samples, heart and liver have 3 samples). Here, accuracy measures the proportion of predicted PSI values that are within 0.2 of the measured PSI values.

Next, we evaluated Splice Ninja’s ability to predict tissue-specific PSI changes by comparing predicted and measured dPSI for every pair of unseen tissues on test chromosomes (Table 2). Overall, predicted and observed dPSI show modest correlation, which increases markedly for significantly differentially-spliced events (|dPSI| ≥ 0.15), and for ES and IR events. Given the difficulty of predicting tissue-specific changes, especially between two unseen tissues, these modest correlations are still indicative of Splice Ninja capturing some tissue-specific splicing mechanisms. Furthermore, we observe a substantial 15–35% overlap between the events Splice Ninja predicts as most differentially spliced and those that actually exhibit the largest dPSI, demonstrating its ability to recover a significant subset of key differentially-spliced events. Additionally, Splice Ninja uncovers many of the most differentially spliced events—regardless of whether dPSI is positive or negative—but struggles to isolate events with minimal splicing changes. Overall, it captures key aspects of tissue-specific splicing, yet its understanding of these mechanisms is still imperfect.

**Table 2:** Performance on predicting dPSI between unseen tissue pairs on test chromosomes. For each pair: number of shared events; Spearman correlation (*ρ*) and Pearson correlation (*r*) for all events and for significant events (|dPSI| ≥ 0.15); and percent overlap between the top/bottom 10% of events by predicted vs. measured signed and absolute dPSI. Predicted and measured PSI values are averaged across samples from a given tissue before computing these metrics.

| Event type | Tissue 1 | Tissue 2 | Num events | $\rho$ | $r$ | Num sig events | Sig events $\rho$ | Sig events $r$ | % top pred diff in top diff | % bottom pred diff in bottom diff | % top pred abs diff in top abs diff | % bottom pred abs diff in bottom abs diff |
| --- | --- | --- | --- | --- | --- | --- | --- | --- | --- | --- | --- | --- |
| ES | Cerebellum | Heart | 36661 | 0.182 | 0.099 | 1132 | 0.237 | 0.228 | 25.777 | 20.731 | 24.768 | 10.611 |
| ES | Cerebellum | Liver | 32667 | 0.180 | 0.163 | 1153 | 0.332 | 0.372 | 23.056 | 20.974 | 22.780 | 9.584 |
| ES | Heart | Liver | 34733 | 0.019 | 0.086 | 683 | 0.268 | 0.266 | 16.211 | 16.873 | 19.349 | 11.258 |
| IR | Cerebellum | Heart | 31358 | 0.230 | 0.221 | 1595 | 0.289 | 0.377 | 24.338 | 25.774 | 11.547 | 7.624 |
| IR | Cerebellum | Liver | 28033 | 0.445 | 0.358 | 2098 | 0.630 | 0.629 | 22.726 | 33.964 | 20.978 | 14.556 |
| IR | Heart | Liver | 30129 | 0.241 | 0.270 | 1268 | 0.368 | 0.444 | 21.614 | 35.989 | 9.728 | 8.333 |
| ALTA | Cerebellum | Heart | 23590 | 0.191 | 0.037 | 224 | 0.006 | -0.058 | 23.485 | 25.943 | 30.818 | 10.343 |
| ALTA | Cerebellum | Liver | 20360 | 0.119 | 0.038 | 212 | -0.010 | 0.010 | 21.365 | 22.397 | 28.929 | 11.248 |
| ALTA | Heart | Liver | 22164 | -0.059 | -0.026 | 169 | -0.098 | -0.110 | 14.576 | 14.395 | 24.052 | 12.951 |
| ALTD | Cerebellum | Heart | 14380 | 0.169 | 0.096 | 135 | 0.240 | 0.210 | 26.634 | 25.313 | 26.356 | 13.561 |
| ALTD | Cerebellum | Liver | 12398 | 0.137 | 0.077 | 165 | 0.177 | 0.203 | 25.020 | 23.406 | 27.764 | 12.026 |
| ALTD | Heart | Liver | 13494 | 0.005 | 0.023 | 108 | 0.092 | 0.187 | 15.864 | 19.051 | 25.204 | 11.193 |

### 5.2 Comparing predictions for unseen tissues with predictions from other models trained on these tissues

We use the benchmarks proposed by Wu et al. [2] to compare Splice Ninja’s performance in unseen tissues with the performance of existing models – Pangolin [1] and TrASPr [2], that are trained on these tissues. SpliceAI is used as an additional baseline to estimate the performance of non-tissue-specific models. The benchmarks concentrate on ES events and to obtain PSI estimates from SpliceAI and Pangolin that only predict splice sites, Wu et al. [2] computed the probability of the 5’ and 3’ splice sites for the alternate exon and averaged them. We directly use the predictions provided by Wu et al. [2] in our analyses. Again, as mentioned in Section 2.2, Wu et al. [2] benchmark TrASPr on two unseen GTEx tissues. However, since we were unable to access the models and predictions used in that analysis, we do not compare TrASPr and Splice Ninja on tissues unseen by both models.

Wu et al. [2] processed GTEx data [17] from six different tissues, including our unseen tissues – cerebellum, liver, and heart, to obtain PSI values for ES events and trained TrASPr on these values. As mentioned in Section 4.2, like Splice Ninja, the other models also use data from chromosomes 1, 3, 5, and 7 in their test sets. Here, we use PSI values and predictions for the cerebellum, liver, and heart published by Wu et al. [2] for ES events from test chromosomes^5^ and compare them to Splice Ninja predictions. Out of the ~2500 events used by Wu et al. [2] for benchmarking, we were able to retrieve splicing event information from VastDB for ~2100 events and we use these events in our results. To get the SF expression levels required for making predictions using Splice Ninja, we obtained median TPM values from GTEx for the cerebellum, liver, and heart and applied the necessary transformations.

Figure 4 shows the overall prediction performance of all models. In general, we observe that Splice Ninja significantly outperforms SpliceAI and has performance that is comparable to Pangolin, another model with a SpliceAI-based architecture. However, it still trails TrASPr, a much larger BERT-based model, and all models perform considerably worse on events with significant PSI shifts between tissue pairs, highlighting the difficulty in predicting tissue-specific splicing events. Importantly, this figure clearly shows the promise of models like Splice Ninja that capture trans-effects. Using a similar model architecture and scale, we are able to achieve prediction performances for unseen tissues that are similar to those obtained using models that are trained on these tissues.

**Fig. 4:**
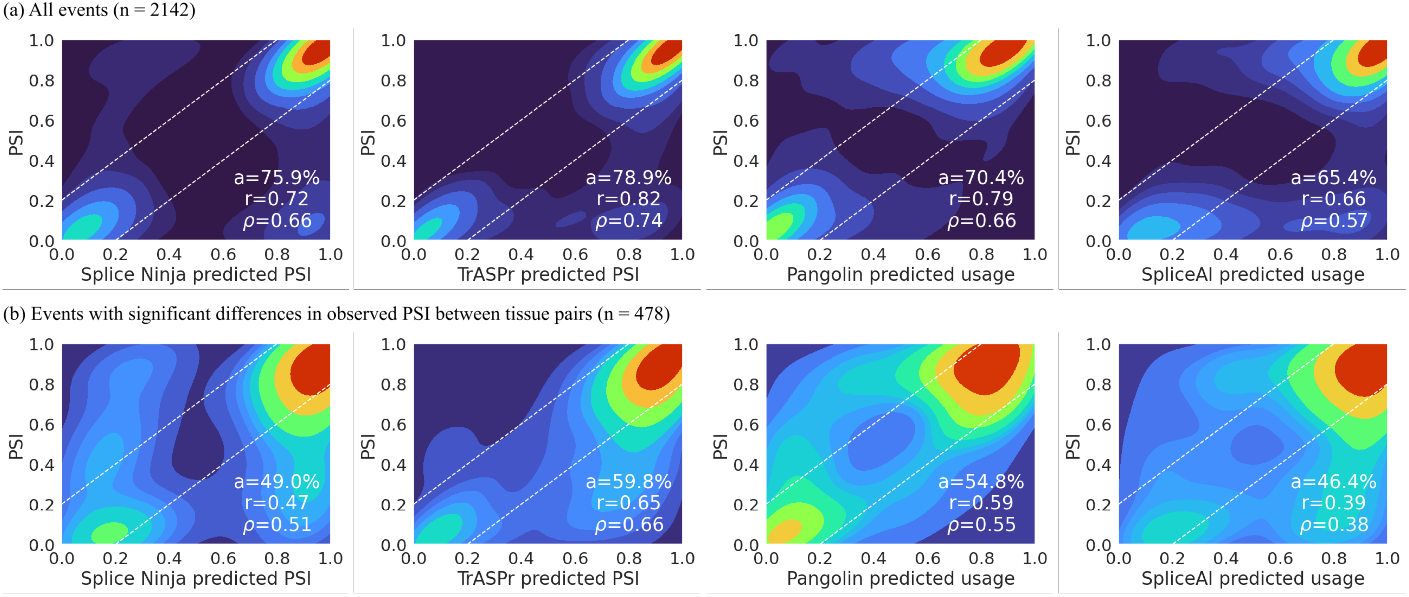
Comparing the performance of Splice Ninja on unseen tissues with the performance of existing methods that are trained on these tissues. SpliceAI is included as a baseline to show the performance of non-tissue-specific model. (a) Results for all splicing events; (b) Results for events exhibiting significant PSI shifts between tissue pairs (dPSI > 0.15). For each model, we report the percentage of predictions within ±0.2 of the measured PSI (“a”), Pearson correlation (*r*), and Spearman correlation (*ρ*). Dashed lines denote ±0.2 deviation from the identity line.

Lastly, we evaluated each model’s ability to flag exons as significantly included (dPSI+) or excluded (dPSI-) (|dPSI| > 0.15) between tissue pairs by framing two binary classification tasks per comparison. Figure 5 reports AUROC and AUPRC for Splice Ninja, Pangolin, and TrASPr. Overall, Splice Ninja’s performance closely matches that of Pangolin—with deficits in a few comparisons—while both models are consistently surpassed by TrASPr.

**Fig. 5:**
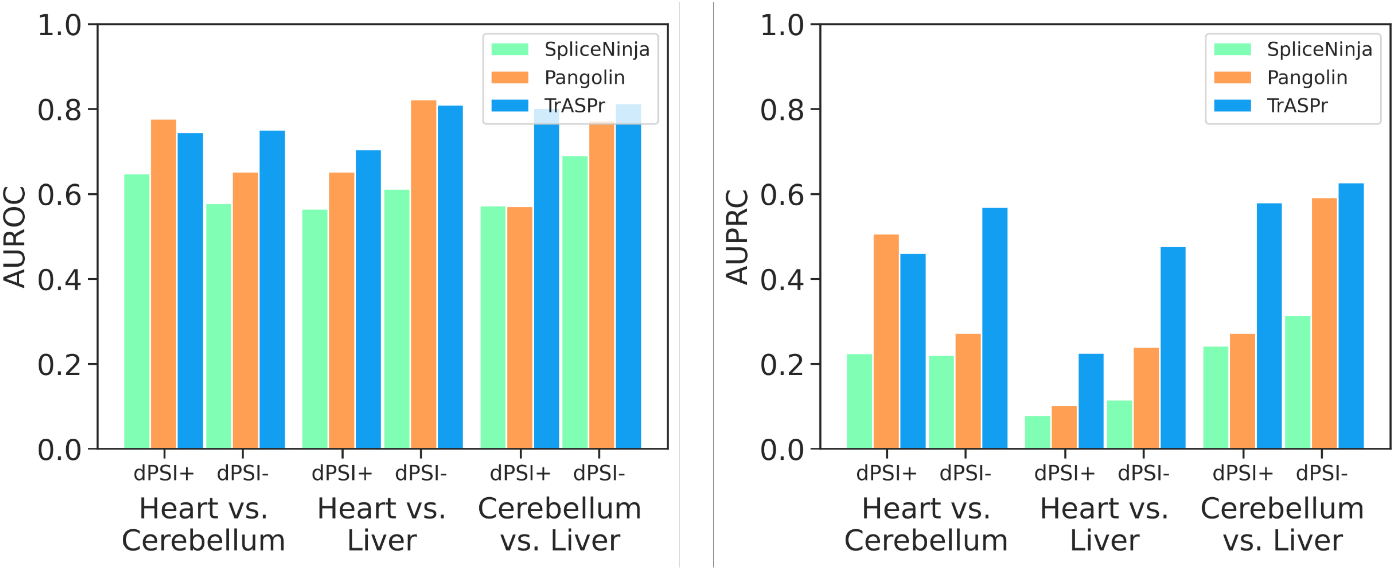
Comparing the performance of Splice Ninja with Pangolin and TrASPr for distinguishing significantly differentially-spliced exons between two tissue pairs (|dPSI > 0.15|) from all other exons. dPSI+ corresponds to differentially included exons and dPSI-corresponds to differentially excluded ones. The panel on the left shows area under the receiver operating characteristic curve (AUROC) and the panel on the right shows area under the precision-recall curve (AUPRC) for different tissue pairs.

## 6 Discussion, limitations, and future work

Being able to predict tissue-specific splicing is important for understanding how splicing mutations have different effects in different tissues. In this work, we introduced Splice Ninja, a deep learning model that addresses several limitations of existing splicing predictors by incorporating SF expression levels into a sequence-based model. Splice Ninja is trained to predict tissue-specific PSI values for splicing events and is designed to make predictions in tissues that were never seen during training.

Our results show that this approach works well. Splice Ninja makes accurate PSI predictions for most events in unseen tissues and is able to identify many tissue-specific splicing changes. In fact, even though it has never seen data from the test tissues, its performance is similar to Pangolin—a model with a similar architecture that is trained directly on those tissues. This suggests that using SF expression as an input provides useful information about tissue-specific splicing regulation.

Nonetheless, there are still some limitations. First, Splice Ninja performs worse on IR events compared to other event types, though it still identifies significantly differentially-retained introns to some extent. This might be because IR events tend to have smaller changes in PSI across tissues, and our current loss function emphasizes larger dPSI values. Changing the loss to give more weight to smaller dPSI values—especially for IR—could help improve performance.

Splice Ninja predictions for the unseen tissues are also worse than those obtained using TrASPr. Even though TrASPr is directly trained on these tissues, adopting some of the techniques used to train it might be beneficial for unseen tissue prediction. TrASPr is trained to predict the dPSI between a pair of tissues, using a contrastive-style approach. This might help it better understand how the splicing environment differs between tissues and could explain its stronger performance in tissue-specific settings. While the TrASPr manuscript reports that it can generalize to unseen cell lines and tissues using RBP expression, we were unable to access the pretrained models or predictions used in their unseen tissue analyses. In general, it would be interesting to try a similar contrastive approach with Splice Ninja and see if it improves performance on unseen tissues.

In conclusion, Splice Ninja shows that adding SF expression to a sequence-based model helps with predicting splicing in tissues that have not been seen during training. While there is room to improve—especially for certain event types and with better training strategies—this modeling approach is a promising step toward building more general splicing predictors.

## A Data-splits for train, validation, and test sets

We use data from the following 129 VastDB samples in the train set:

Adipose_b, Adipose_c, Adipose_d, Adrenal_b, Adrenal_c, Amnion, Astro-cytes, Bladder_a, Bone_marrow_a, Bone_marrow_b, Bone_marrow_c, Brain_Endoth, Breast_a, Breast_Epith_a, Chorion, CL_293T, CL_EndoBH1C_a, CL_Gm12878, CL_HeLa, CL_K562, CL_LP1, CL_MB231, CL_MCF7, CL_PNT2, CL_SHSY5Y_noRA, CL_SHSY5Y_RA, Colon_b, Cortex, Decidua, Embr_2C_a_A, Embr_2C_a_B, Embr_4C_a_A, Embr_4C_a_B, Embr_4C_a_C, Embr_8C_a_A, Embr_8C_a_B, Embr_8C_a_C, Embr_8C_a_D, Embr_Cortex_13_17wpc, Embr_Forebrain_9_12wpc, Embr_Forebrain_St13_14, Embr_Forebrain_St17_20, Embr_Forebrain_St22_23, Embr_ICM_a_A, Embr_Morula_a_A, Embr_Morula_a_B, Embr_TE_a_A, Embr_TE_a_B, EndomStromCells, EndothCells, EpithelialCells, ESC_H1_a, ESC_H1_b, ESC_H1_c, ESC_H1_d, ESC_H9_a, ESC_H9_b, Fibroblasts, Frontal_Gyrus_old, Frontal_Gyrus_young, GLS_cells, HFDPC, HMEpC_a, iPS_a, iPS_b, Kidney_d, Kidney_e, Kidney_f, Lung_b, Lung_e, Lung_f, Lymph_node_b, Lymph_node_c, Melanocytes, Microglia, MonoNucCells, MSC, Muscle_b, Muscle_d, Muscle_e, NCC_Cranial_c, NCC_default_b, NCC_Enteric_b, NCC_Melano_biased_b, Neuroblastoma, Neurons, Neurons_Cortex_KCl_0h, Neurons_Cortex_KCl_1h, Neu-rons_Cortex_KCl_6h, NPC_a, NPC_b, Oligodendrocytes, Oocyte_a_A, Ovary_a, Ovary_b, Pancreas_Acinar_Old, Pancreas_Acinar_Young, Pancreas_Alpha_Old, Pancreas_Alpha_Young, Pancreas_Beta_Old, Pancreas_Beta_Young, Pan-creas_Duct_Old, Pancreas_Duct_Young, Placenta_a, Placenta_b, Placenta_c, Placenta_Epith, Prostate_b, Prostate_c, Prostate_d, Retina_a, Small_intestine, Spleen_a, Spleen_b, Stomach_a, Stomach_b, Sup_Temporal_Gyrus, Testis_a, Testis_b, Testis_c, Thymus_a, Thymus_b, Thyroid_b, Thyroid_c, Thyroid_d, WBC_d, WBC_e, Whole_Brain_b, Zygote_a_A.

The following 8 samples are used in the validation set:

Colon_sigmoid, Colon_transverse, Retina_macular, Retina_peripheral, Skin_a, Skin_b, Skin_c, Skin_d

Finally, the following 8 samples are used in the test set:

Cerebellum_a, Cerebellum_c, Liver_a, Liver_b, Liver_c, Heart_a, Heart_b, Heart_c

## Supplementary Figures

**Fig. S1:**
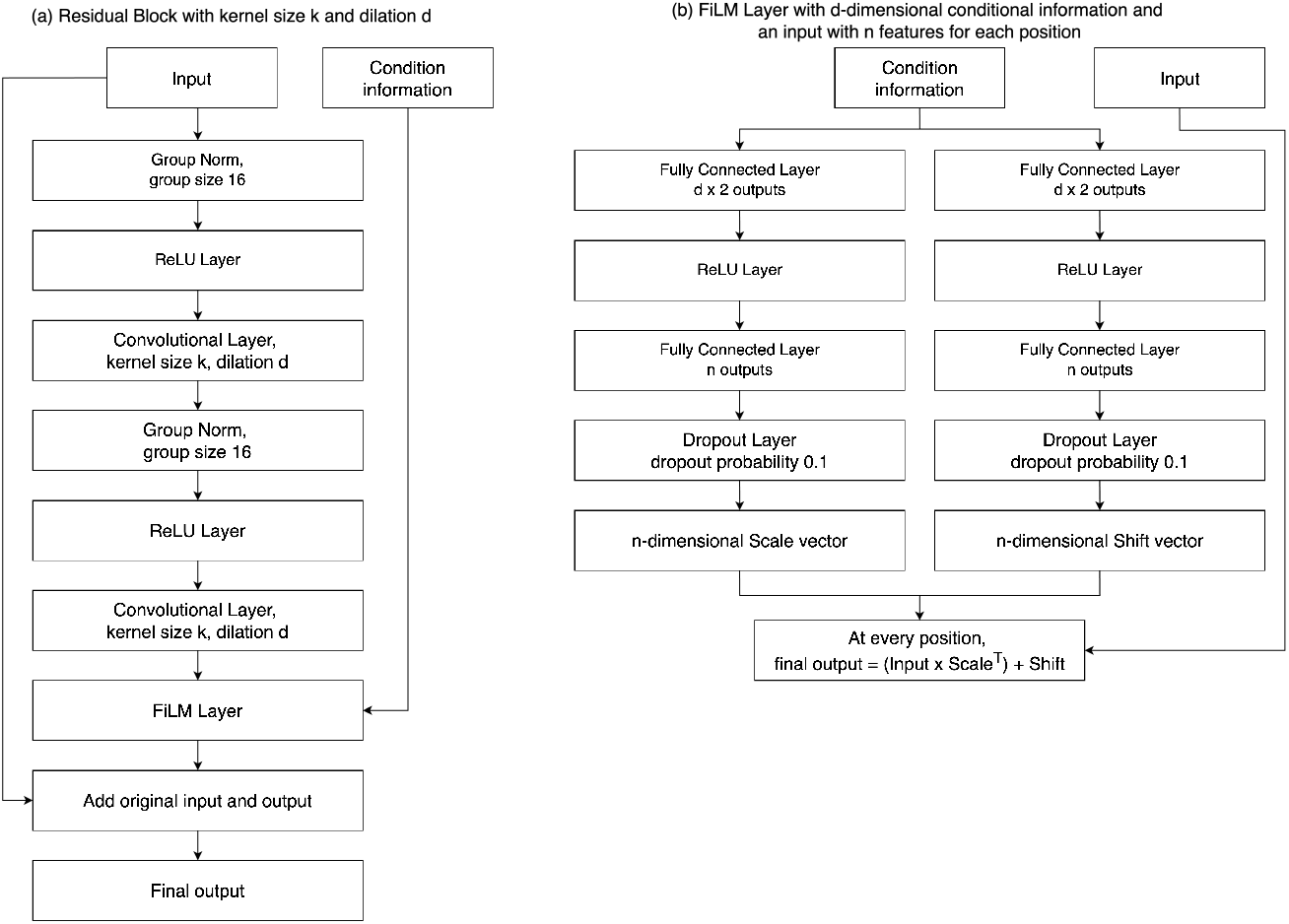
Detailed illustrations of the modules used in Splice Ninja. Panel (a) shows the structure of the residual block we use and panel (b) details the Feature-wise Linear Modulation or FiLM layers we use for incorporating conditioning information while making predictions.

## Footnotes

1 In all our analyses, instead of using percentages, we use PSI values in the range [0, 1].

2 For splicing events exceeding 10,000 bp, we iteratively trim central intronic (or exonic for IR events) regions while preserving 500 bp at each end, then extend to 10,000 bp with adjacent genomic context.

3 Reads per kilobase for gene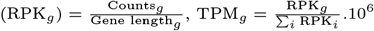

4 An epoch is defined as seeing as many PSI values as in the train set.

5 https://github.com/AstroSign/TrASPr_model/blob/main/plot_script/data/GTEx_data.tsv

